# Mathematical model of tumour spheroid experiments with real-time cell cycle imaging

**DOI:** 10.1101/2020.12.06.413856

**Authors:** Wang Jin, Loredana Spoerri, Nikolas K. Haass, Matthew J. Simpson

**Affiliations:** School of Mathematical Sciences, Queensland University of Technology, Brisbane, Queensland 4001, Australia; The University of Queensland, The University of Queensland, Diamantina Institute, Translational Research Institute, Brisbane, Australia

**Keywords:** Mathematical modelling, Tumour spheroid, Cell cycle, Cancer, Moving boundary problem

## Abstract

Three-dimensional (3D) *in vitro* tumour spheroid experiments are an important tool for studying cancer progression and potential drug therapies. Standard experiments involve growing and imaging spheroids to explore how different experimental conditions lead to different rates of spheroid growth. These kinds of experiments, however, do not reveal any information about the spatial distribution of the cell cycle within the expanding spheroid. Since 2008, a new experimental technology called fluorescent ubiquitination-based cell cycle indicator (FUCCI), has enabled real time *in situ* visualisation of the cell cycle progression. FUCCI labelling involves cells in G1 phase of the cell cycle fluorescing red, and cells in the S/G2/M phase of the cell cycle fluorescing green. Experimental observations of 3D tumour spheroids with FUCCI labelling reveal significant intratumoural structure, as the cell cycle status can vary with location. Although many mathematical models of tumour spheroid growth have been developed, none of the existing mathematical models are designed to interpret experimental observations with FUCCI labelling. In this work we extend the mathematical framework originally proposed by Ward and King (1997) to develop a new mathematical model of FUCCI-labelled tumour spheroid growth. The mathematical model treats the spheroid as being composed of three subpopulations: (i) living cells in G1 phase that fluoresce red; (ii) living cells in S/G2/M phase that fluoresce green; and, (iii) dead cells that do not fluoresce. We assume that the rates at which cells pass through different phases of the cell cycle, and the rate of cell death, depend upon the local oxygen concentration in the spheroid. Parameterising the new mathematical model using experimental measurements of cell cycle transition times, we show that the model can capture important experimental observations that cannot be addressed using previous mathematical models. Further, we show that the mathematical model can be used to quantitatively mimic the action of anti-mitotic drugs applied to the spheroid. All software required to solve the nonlinear moving boundary problem associated with the new mathematical model are available on [GitHub](https://github.com/wang-jin-mathbio/jin2021).

## 1 Introduction

*In vitro* three-dimensional (3D) tumour spheroids have been widely used to study cancer progression and putative therapies (Kunz-Schughart et al., 1998; Nath and Devi, 2016; Santini and Rainaldi, 1999; Spoerri et al., 2017). The generation and analysis of tumour spheroids have been documented in the literature for approximately fifty years (Sutherland et al., 1971; Greenspan, 1972). Traditional spheroid experiments involve growing populations of cancer cells in 3D culture, and visualising spheroid growth and invasion by taking phase contrast images (Beaumont et al., 2014). As an illustrative example, the image in Figure 1(a) shows a spheroid generated from C8161 melanoma cells (Nath and Devi, 2016). This type of experiment can be used to measure the expansion of the tumour spheroid over time. In some experiments the formation of a necrotic core within the spheroid can be examined using markers of cell death (Chan et al., 2003; Maeda et al. 2014), and fluorescent live/dead assays (e.g. Calcein-AM/Ethidium bromide or DRAQ7) have been used to assess drug cytotoxicity (Smalley et al., 2008; Kienzle et al., 2017). This standard experimental protocol provides very little insight into cell cycle behaviour and spatio-temporal distribution patterns of proliferating cells, within the growing spheroid.

**Figure 1:**
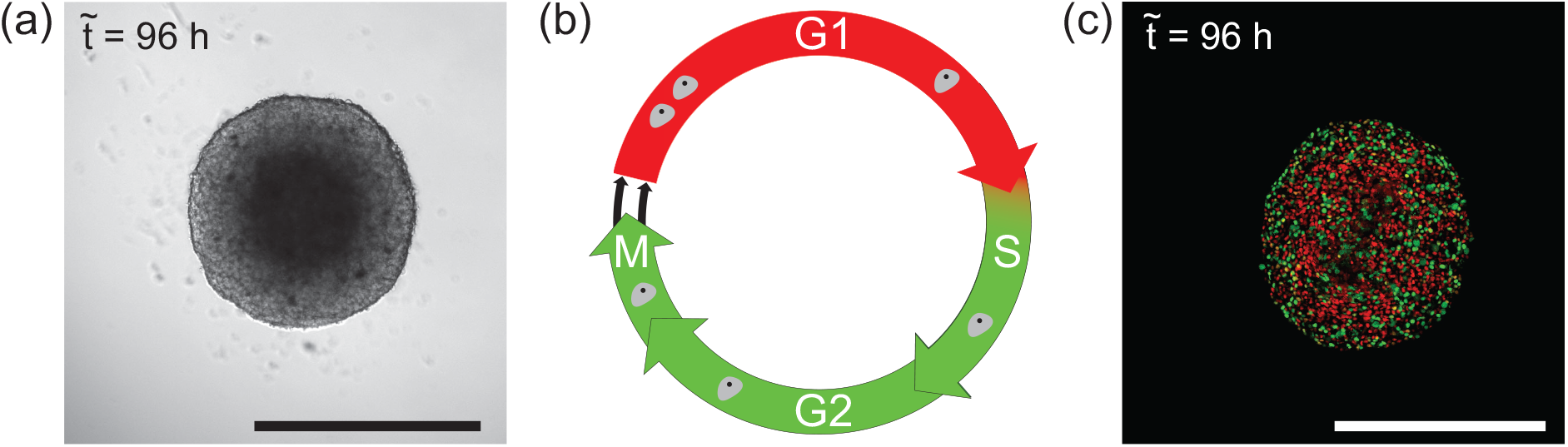
Motivating images and cell cycle schematic. (a) Phase contrast image of a 3D melanoma spheroid with the C8161 cell line grown on agar after 96 h. The scale bar corresponds to 600 *μ*m. (b) Schematic of the eukaryotic cell cycle indicating the transition between different phases together with the red and green fluorescence colours associated with FUCCI labelling. For the purpose of this study, we neglect double-positivity (yellow) in early S phase. (c) Confocal microscopy image of a 3D melanoma spheroid after 96 h, using FUCCI-transduced C8161 melanoma cells. The scale bar corresponds to 400 *μ*m.

Spheroids mimic physiological tumour features, such as cell cycle behaviour, which is responsible for tumour cell proliferation and thus tumour growth, but also invasion, cell-cell and cell-matrix interactions, molecule diffusion, oxygen/nutrient gradients (with a hypoxic zone and a central necrosis) as well as drug sensitivity (Beaumont et al., 2014; Smalley et al., 2008; Spoerri et al., 2017). The cell cycle consists of a sequence of four distinct phases: gap 1 (G1), synthesis (S), gap 2 (G2), and the mitotic (M) phase, as illustrated in Figure 1(b). The G1, S, and G2 phases are referred to collectively as *interphase*, which involves cell growth and preparation for division. After interphase, the cell enters the mitotic phase and divides into two daughter cells. Since 2008 (Sakaue-Sawano et al., 2008), a new experimental technique called *fluorescent ubiquitination-based cell cycle indicator* (FUCCI) has enabled real time *in situ* visualisation of the cell cycle progression from G1 to S/G2/M in individual cells. The FUCCI system consists of two fluorescent probes which emit red fluorescence when the cell is in the G1 phase, or green fluorescence when the cell is in S/G2/M phase (Sakaue-Sawano et al., 2008), as shown schematically in Figure 1(b). Before the development of FUCCI, it was difficult to examine the cell cycle dynamics of individual cells within a growing population. The spheroids in Figure 1 are constructed with the same melanoma cell line: C8161 (Figure 1(a)) and C8161 expressing FUCCI (Figure 1(c)). The key difference is that FUCCI reveals more detailed information about the cell cycle status at different locations within the growing spheroid. In Figure 1(c) we see that the spheroid periphery has an even distribution of red and green cells, indicating that they are in active cell cycle, i.e. proliferating. In contrast, cells in the spheroid centre are predominantly red, indicating G1-arrest (Haass et al., 2014). This information, revealed by FUCCI, allows us to visualise and measure both the time evolution of the spheroid size and spatial patterns of cell cycle behaviour within the spheroid (Haass et al., 2014). Understanding the biology of cell cycle behaviour is important, as cell cycle phase-specific drug resistance has been described a an escape mechanism of cancer cells (Beaumont et al., 2016; Haass and Gabrielli, 2017).

Several mathematical models have been developed to describe experimental tumour spheroids since the 1970s (Greenspan, 1972; McElwain et al. 1978; McElwain et al. 1979; Norton et al., 1976). The seminal mathematical model by Greenspan (1972) idealises a tumour spheroid as a compound sphere: the inner-most region is a necrotic core containing dead cells; the intermediate spherical shell contains living, non-proliferative, or *quiescent* cells; and the outer-most spherical shell contains live, freely–proliferating cells. A key assumption in Greenspan’s model is that cell proliferation is limited by the availability of a nutrient, such as oxygen. Greenspan (1972) assumes that oxygen diffuses into the spheroid from the free, growing surface. Living cells are assumed to consume oxygen, and those cells die when there is insufficient oxygen available locally. A further assumption is that dead cells produce a chemical inhibitor, such as metabolites, that diffuse from the necrotic core towards the growing surface of the spheroid. Assembling these assumptions together in terms of a spherically-symmetric conservation statement, Greenspan’s model can be written as a system of coupled partial differential equations (PDE). The solution of these PDEs describe the time evolution of the spheroid in terms of: (i) the radius of the necrotic core; (ii) the thickness of the spherical shell containing quiescent cells; and, (iii) the outer radius of the spheroid (Greenspan, 1972).

Since Greenspan’s model was first proposed, a number of partially refined models have been introduced (e.g. Deakin, 1975; Pettet et al., 2001; Ward and King, 1997; Flegg and Nataraj, 2019; Sarapata and de Pillis, 2014). An important contribution is the work of Ward and King (1997) who proposed a mathematical model describing a spherically-symmetric population of cells that is composed of a subpopulation of live cells and a subpopulation of dead cells. This model also considers the spatial and temporal distribution of some nutrient, which is again taken to be oxygen. Ward and King (1997) assume that living cells proliferate with a rate that depends on the local oxygen concentration, and that the act of proliferation consumes oxygen. Further, living cells are assumed to die, and death rate is also a function of local oxygen concentration. All cells in the spheroid move with a local velocity field that is created by net volume gain as a result of cell proliferation and cell death, since they make the reasonable assumption that the volume of a dead cell is less than the volume of a living cell. Together, these assumptions lead to a novel moving boundary PDE model. The solution of this moving boundary problem can, for certain choices of parameters, lead to the formation of a necrotic core in the growing spheroid (Ward and King, 1997). A key difference between the models of Greenspan (1972) and Ward and King (1997) is that the former defines the interfaces between the necrotic core and the quiescent zone, and between the quiescent zone and the freely proliferating zone to be perfectly sharp well–defined interfaces, while the latter is continuous, and the internal boundaries are less well–defined. Other approaches to modelling tumour spheroids, including multiphase models, have also been developed (Breward et al., 2002; Byrne and Preziosi, 2003; Chaplain et al., 2006; Collis et al., 2016; Enderling and Chaplain, 2014; Landman and Please, 2001; Lewin et al., 2020; Spill et al., 2016), however none of these existing models are suitable for interpreting experimental data with FUCCI labelling.

In this study we present a mathematical model of *in vitro* tumour spheroids that is an extension of Ward and King’s modelling framework. The key feature of our model is that we explicitly incorporate the transition between different phases of the cell cycle so that the model can be applied to tumour spheroids with FUCCI labelling. Our model considers the spheroid to be composed of three subpopulations: (i) living cells in G1 phase that fluoresce red; (ii) living cells in S/G2/M phase that fluoresce green; and, (iii) non–florescent dead cells. We assume that the rates at which cells progress through the cell cycle depend upon the local oxygen concentration, and that the rate of cell death also depends on the local oxygen concentration in the spheroid. We partially parameterise the new mathematical model using existing experimental measurements of cell cycle transition times, and then, using additional carefully chosen parameter values, we show that the model captures important experimental observations that cannot be observed in standard experiments without FUCCI labelling. We conclude by exploring how the model can be used to simulate the application of anti-mitotic drugs.

## 2 Experimental motivation

Our modelling work is motivated by experimental observations of tumor spheroids with FUCCI labelling, such as the image in Figure 1(c). In Figure 2 we present snapshots from a time-lapse movie of FUCCI-labelled spheroid growth, again with FUCCI-C8161 melanoma cells. At early time, 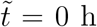, we see that the spheroid is relatively homogeneous, and composed of well–mixed red and green fluorescing cells with no obvious spatial patterns. At intermediate time, 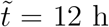, the spheroid has expanded in radius, and the central region of the spheroid is composed mostly of cells in G1 phase (red), whereas the outer shell of the spheroid contains well–mixed red and green fluorescing cells. At later time, 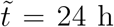, we see the spheroid continues to expand in size. In this case the outer spherical shell contains well–mixed red and green fluorescing cells, an intermediate spherical shell is composed of predominantly red fluorescing cells, whereas the absence of florescence in the central region indicates the formation of a necrotic core. These images show the importance of FUCCI labelling as this information indicates that we have a strong correlation between the spatial position and the cell cycle status. In a spheroid without FUCCI labelling, such as in the image in Figure 1(a), it is not possible to see this level of detail. Further, this tight coupling between location and cell cycle status illustrates the biological importance of working in a 3D geometry, as it is largely absent in two-dimensional (2D) experiments where all cells in the population have access to oxygen since these experiments involve a thin monolayer of cells (Haass et al., 2014; Vittadello et al., 2018; Simpson et al., 2020).

**Figure 2:**
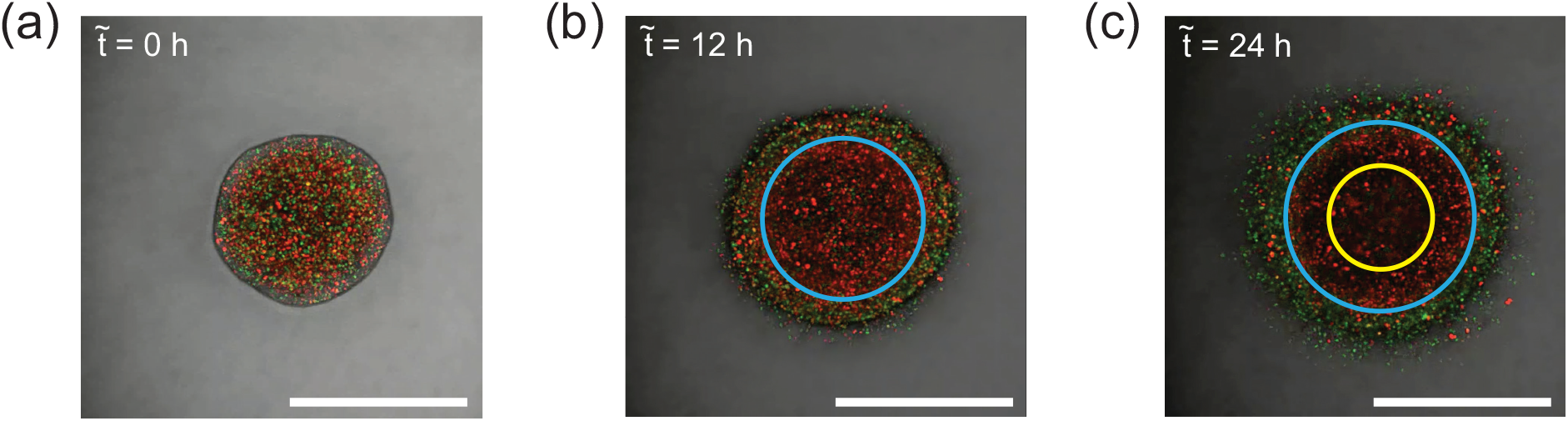
*In vitro* tumour spheroid generated with FUCCI-C8161 melanoma cells. (a)-(c) Still images from a time lapse movie over 24 h, with the times as indicated. (a) shows the red and green fluorescing cells relatively evenly distributed throughout the spheroid. (b) shows a mixture of red and green fluorescing cells at the surface of the spheroid, whereas in the centre of the spheroid we see predominantly red fluorescing cells. (c) shows the inner-most region is largely free of fluorescing cells; the intermediate region contains mostly red fluorescing cells; and the outermost region contains a mixture of red and green fluorescing cells. Blue and yellow concentric circles are superimposed on (b) and (c) to show the approximate boundaries between these zones. The scale bars correspond to 500 *μ*m.

We hypothesize that the simplest explanation of the spatial patterns in cell cycle status in Figure 2 is caused by differences in local nutrient availability. As in the case of Greenspan (1972) and Ward and King (1997), we make the assumption that the key nutrient of interest is oxygen. Furthermore, if we assume that oxygen diffuses into the spheroid from the free surface, we expect that cells near the spheroid surface will have access to sufficient oxygen to proliferate freely. Since cells consume oxygen to survive and proliferate, we anticipate that the oxygen concentration will decrease with depth from the spheroid surface. Accordingly, at some depth, we expect that there could be sufficient oxygen to enable cell survival, but insufficient to allow cells to enter the cell cycle. This would explain the formation of a living, quiescent, G1-arrested population of cells at intermediate depths within the sphere. At locations deeper within the spheroid still, where oxygen concentration would be smallest, the local oxygen concentration would be insufficient to support living cells, giving rise to a necrotic region.

To translate these biological observations and assumptions into a mathematical model, we first present a schematic of the cell cycle, coupled with FUCCI labelling, in Figure 3. The schematic in Figure 3(a), for plentiful oxygen availability, indicates that the cell cycle proceeds freely, and cells in G1 phase (red) enter the cell cycle at a particular rate and proceed through the S/G2/M phase (green) before dividing, at another particular rate, into two daughter cells in G1 phase (red). In this schematic we also allow all living cells to undergo some intrinsic death process. The schematic in Figure 3(b), for intermediate oxygen availability, indicates that the cell cycle is interrupted by the reduced oxygen availability. Under these intermediate oxygen conditions cells in G1 phase (red) are not able to enter the cell cycle because of the reduced oxygen availability. However, those cells already in S/G2/M phase (green) are able to continue through the cell cycle before dividing, to produce two daughter cells in G1 phase (red). The schematic in Figure 3(c), for further reduced oxygen availability, indicates that both the transition from G1 phase (red) to S/G2/M phase (green), and the transition from S/G2/M phase (green) to G1 phase (red) could be interrupted because of the reduced oxygen availability. In this case we expect that the dominant mechanism affecting the population dynamics is localised cell death.

**Figure 3:**
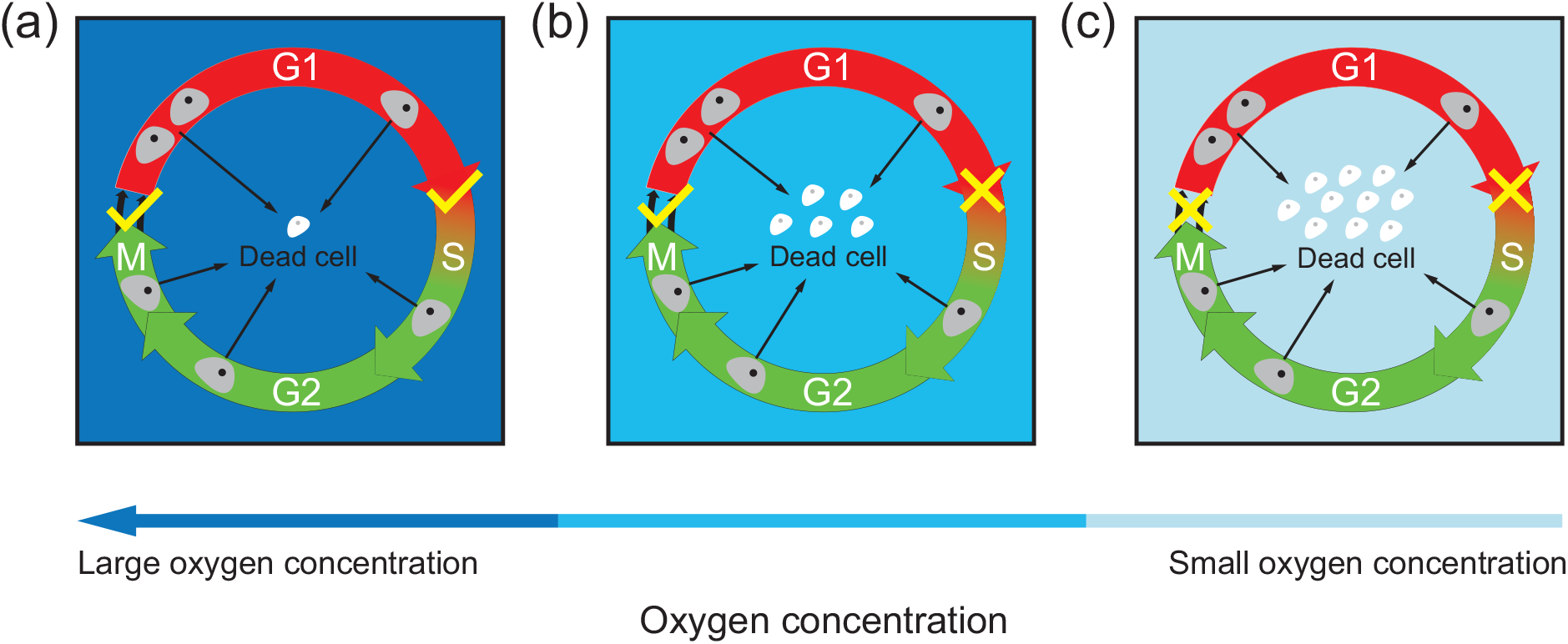
Schematic of oxygen-dependent transitions in the cell cycle with FUCCI la-belling. (a) When oxygen is abundant, cells in G1 phase (red) transition to S/G2/M phase (green), and cells in S/G2/M phase (green) divide into two daughter cells in G1 phase (red). (b) When oxygen availability is reduced, cells in S/G2/M phase (green) continue through the cell cycle and divide into two daughter cells in G1 phase (red), however, those cells in G1 phase (red) do not have access to sufficient oxygen to proceed through the cell cycle. (c) When oxygen availability is sufficiently reduced both the transitions from G1 to S/G2/M, and from S/G2/M to G1 are interrupted.

Based on the experimental observations in Figure 2 (Haass et al., 2014), and the schematic in Figure 3, we now construct a mathematical model that can be used to interpret experimental data describing 3D tumour spheroids with FUCCI by extending the framework originally proposed by Ward and King (1997).

## 3 Model formulation

In the initial presentation of the model we consider all quantities to be dimensional, and denoted using a tilde. Later, we will simplify the model by nondimensionalising all dependent and independent variables. All nondimensional quantities are denoted using regular variables, without the tilde. We take 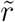 to be the volume fraction of red fluorescing cells (G1 phase) and 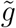 to be the volume green fluorescing cells (S/G2/M phase), so that 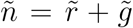 is the volume fraction of living cells, and we suppose that 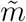 is the volume fraction of dead cells. Mass conservation of the 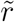 and 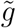 subpopulations is governed by the transitions between different phases of the cell cycle, cell death, and transport by a local velocity field, 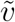, giving

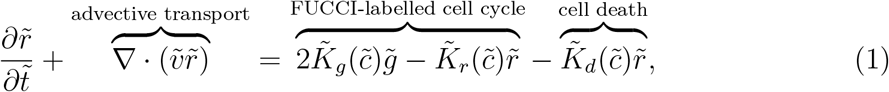

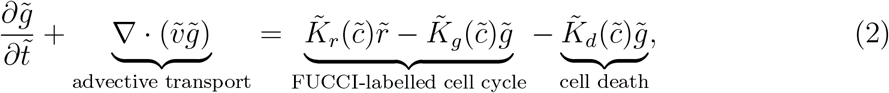

where 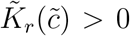 is the rate of transition from G1 to S/G2/M phase, 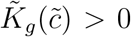 is the rate of transition from S/G2/M to G1 phase, and 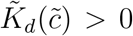 is the rate of cell death. The key feature of our model is that each of these rates depends upon the local oxygen concentration, 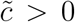. The source terms in Equations (1)–(2) have a relatively simple interpretation. The negative source terms in Equation (1) represent the local loss of cells in G1 phase owing to cell death and the progression through the cell cycle. The positive source term in Equation (2) represent the increase in cells in G2 phase as a result of cells in G1 phase progressing through the cell cycle. The negative source terms in Equation (2) represent the local decrease cells in G2 phase owing to cell death and the progression through the cell cycle. The positive source term in Equation (1) represents the local increase in cells in G1 phase due to the cell cycle, and the factor of two represents the fact that each cell passing from S/G2/M phase to G1 phase produces two daughter cells in G1 phase (Vittadello et al., 2018; Simpson et al., 2020). A useful feature of this nutrient–dependent cell proliferation and nutrient–dependent cell death framework is that it avoids the need for specifying a carrying capacity density as a model parameter, such as in the ubiquitous logistic growth model (Maini et al., 2004a; Maini et al., 2004b; Jin et al., 2017; Jin et al., 2019).

The spatial and temporal distribution of the fraction of dead cells can also be described in terms of a PDE,

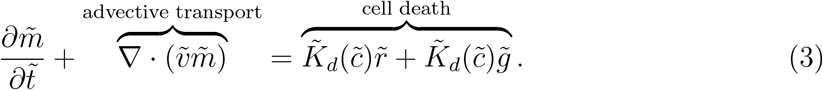

Following Ward and King (1997) we assume the local velocity is associated with net volume gain associated with cell proliferation and cell death,

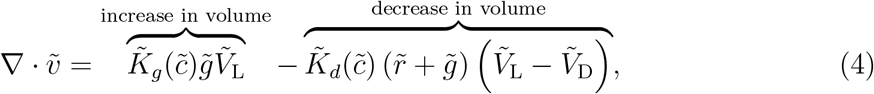

where 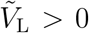 and 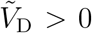 are the volumes of single living and dead cells, respectively. The first term on the right of Equation (4) represents the increase in volume by mitosis, whereas the second term on the right of Equation (4) is the net decrease in volume by cell death. Here we assume that the volume of all living cells is the same, regardless of whether they are in G1 or S/G2/M phase.

Again, following Ward and King (1997) we assume that oxygen is transported with the local velocity as well as through molecular diffusion, with diffusivity 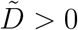, and that oxygen is consumed by living cells, giving rise to

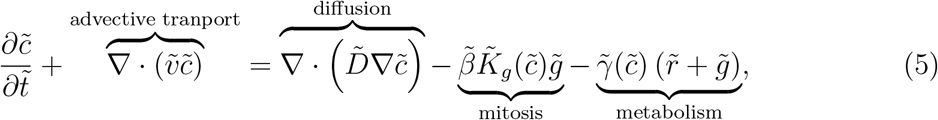

here 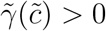 is the rate at which living cells consume oxygen through regular metabolic activity and 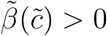 is the rate at which oxygen is consumed during mitosis.

In this model the kinetic rate functions 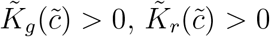, and 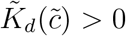 encode critical information about how the rate of transitions between subpopulation 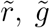, and 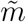 depend upon local oxygen concentration. Based on the experimental observations discussed in Section 2, we suppose these rate functions can be expressed in terms of Hill functions

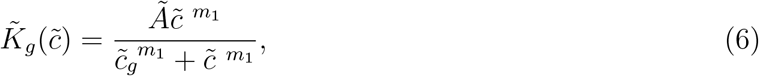

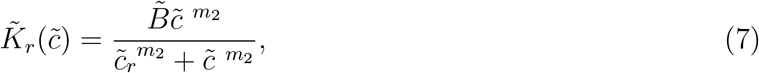

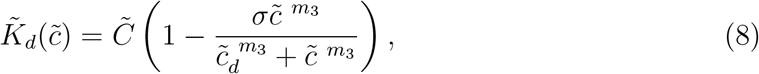

where *Ã* > 0 is the maximum rate at which cells transition from S/G2/M (green) phase to G1(red) phase; 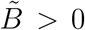 is the maximum rate at which cell transition from G1 (red) phase to S/G2/M (green) phase; 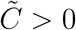 is the maximum rate at which cells die; *m*_1_ > 0, *m*_2_ > 0, and *m*_3_ > 0 are exponents in the Hill functions that govern the steepness about the critical oxygen concentrations 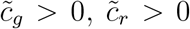, and 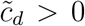, respectively. To ensure 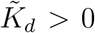 we require 0 ≤ *σ* ≤ 1. Our choice of working with Hill functions in Equations (6)–(30) is not unique, and other functional forms could be used. However, here we choose to work with Hill functions because this is consistent with the original model of Ward and King (1997).

Some further assumptions are required to completely specify the mathematical model. First, assuming there is no voids in the spheroids leads to

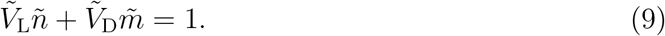

Second, we make the usual assumption that the spheroid is spherically–symmetric so that the two independent variables are the radial position, 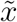, and time, 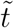. To close the model we specify initial conditions and boundary conditions. Initially, we assume that the spheroid is a mixture of living cells only, leading to

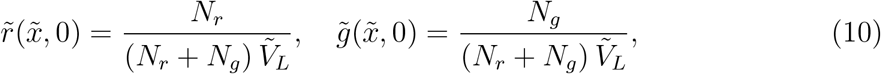

where *N*_*r*_ > 0 and *N*_*g*_ > 0 are the number of red and green cells at 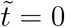. Therefore, the initial radius is

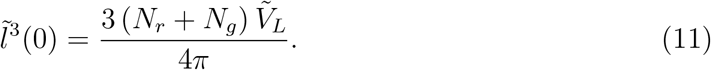

For oxygen, we assume the initial spheroid is sufficiently small that we have

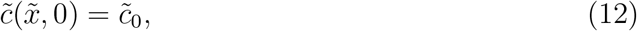

where 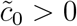 is the external oxygen concentration.

We impose the following boundary conditions by assuming that the time rate of change of the spheroid radius, 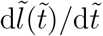, is given by the local velocity at the spheroid surface, and that the oxygen concentration at the spheroid surface is given by the external oxygen concentration,

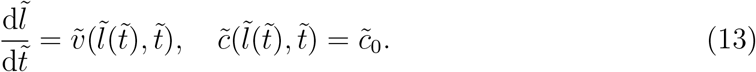

Lastly, we have a symmetry condition at the centre of the spheroid, 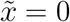,

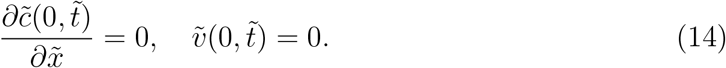

To solve the dimensional model we must specify values of 15 free parameters, (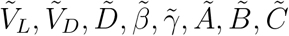 and four quantities relating to initial data, 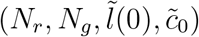. Given the large parameter space for this model we simplify the model through nondimensionalisation.

### 3.1 Nondimensionalisation

Introducing 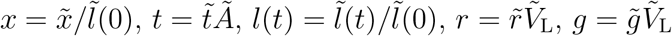, and 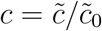 to arrive at a simplified nondimensional model,

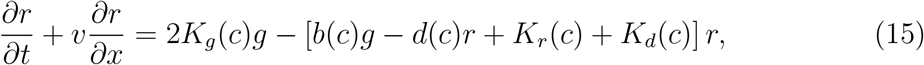

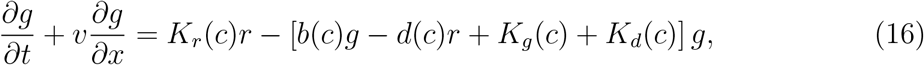

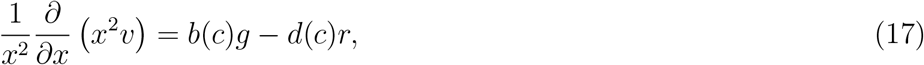

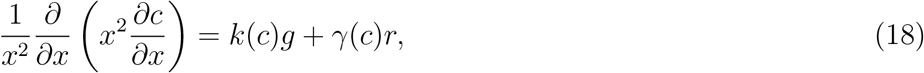

on 0 < *x* < *l*(*t*) and *t* > 0, where *b*(*c*) = *K*_*r*_(*c*) − *d*(*c*), *d*(*c*) = (1 − *δ*)*K*_*d*_(*c*), *k*(*c*) = *β K*_*g*_(*c*) + *γ*(*c*), *δ* = *V*_D_*/V*_L_, 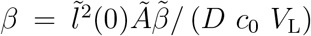, and 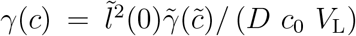. To arrive at Equation (28) from Equation (5), we make the usual quasi-steady-state assumption for oxygen (Greenspan, 1972; Ward and King, 1997).

The nondimensional kinetic rate functions simplify to

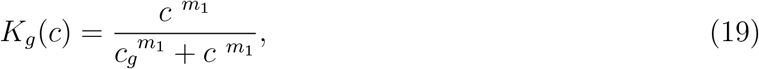

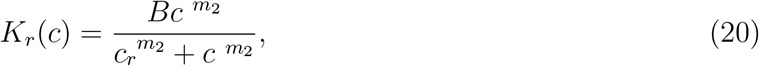

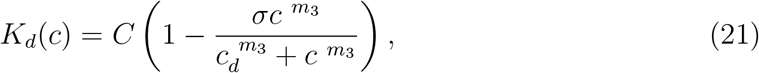

where 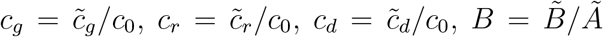, and 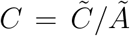. The initial and boundary conditions are given by

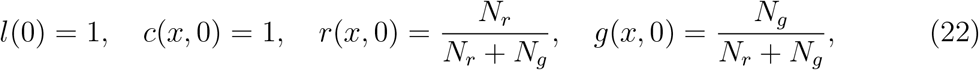

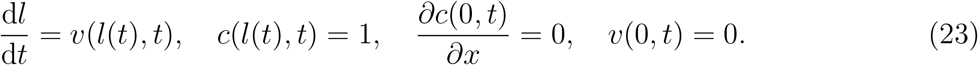

Together, Equations (15)–(28) form a nonlinear moving boundary problem on 0 < *x* < *l*(*t*). We solve the model numerically by first transforming to a fixed domain. The terms associated with the advective transport are approximated using an upstream difference approximation, whereas terms associated with diffusive transport are approximated using a central difference approximation (Simpson et al., 2005). The resulting system of nonlinear evolution equations are integrated through time using an implicit Euler approximation, leading to a system of nonlinear algebraic equations are solved using the Newton-Raphson method. Full details of the numerical method are outlined in the Supplementary Material document, software written in MATLAB are available on [GitHub](https://github.com/wang-jin-mathbio/jin2021).

In our experiments, such those illustrated in Figure 1(c) and Figure 3(a), the initial spheroid radius 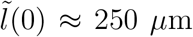. The characteristic timescale is related to the rate parameter *Ã*, which we can estimate by considering experimental data reported by Haass et al. (2014) who make careful measurements of the duration that C8161 melanoma cells fluoresce red and green in 2D culture where oxygen is plentiful. Data in Figure 4(a) show estimates of the duration of time C8161 cells spend in G1 (red) and S/G2/M (green) phase for 20 individual cells. Taking the reciprocal of each recorded duration, we obtain estimates of *Ã* and 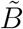. Data in Figure 4(b) suggest that 0.08 < *Ã* < 0.25 /h and 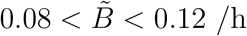. For simplicity we take a representative estimate of *Ã* ≈ 0.1 /h, which means that each unit duration of time in the nondimensional model corresponds to 10 h in the experiment. Given our estimates of *Ã* and 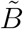, we also estimate 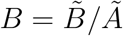, as shown in Figure 4(c), for which we can take a representative estimate of *B* = 1.5. Cell cycle duration estimates differ from cell line to cell line (Haass et al., 2014). Therefore, given this kind of duration data, we could follow a similar procedure to arrive at different estimates of *Ã* and 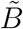 for different cell lines as appropriate.

**Figure 4:**
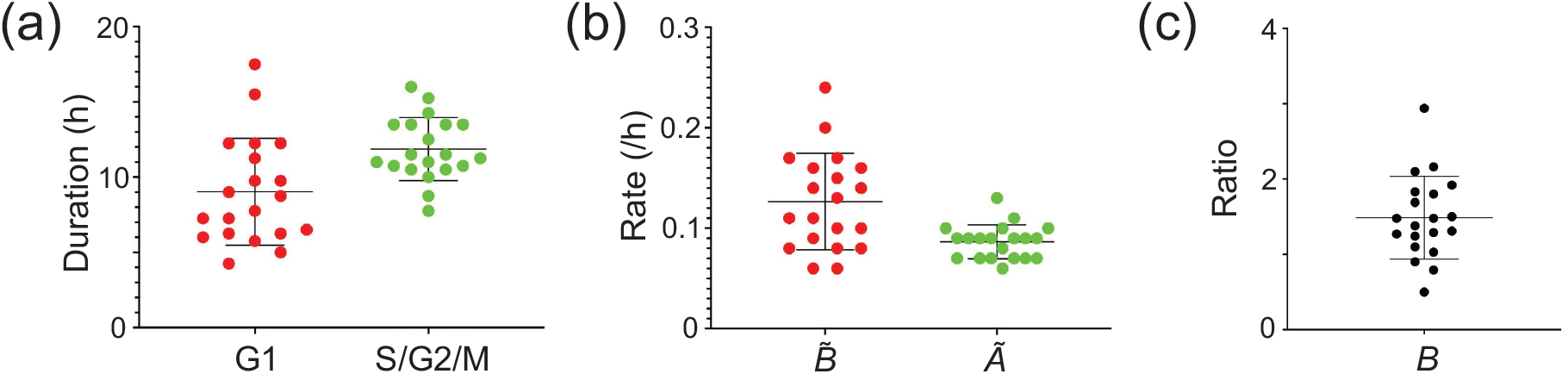
Experimental measurements of cell cycle transition times for FUCCI-C8161 cells without oxygen limitation. (a) Durations of time spent in G1 (red) and S/G2/M (green) phases for twenty FUCCI-C8161 melanoma cells. (b) Estimates of *Ã*, the rate at which cells in S/G2/M (green) go through mitosis and transit into G1 (red), and estimates of 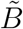, the rate at which cells in G1 (red) enter the cell cycle and transit into S/G2/M (green) phase. (c) Estimates of 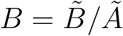.

## 4 Results and discussion

In this section we explore the capability of the mathematical model to describe important features of tumour spheroids with FUCCI labelling. We conduct this exploration by investigating appropriate numerical solutions of Equations (15)–(28). Our aim in this exploration is not to undertake a detailed parameter estimation exercise, but rather to explore whether the new mathematical model, Equations (15)–(28), can replicate key biological features of FUCCI-labelled spheroids. At first we explore a series of *control* experiments by using the mathematical model to replicate various experimental observations, across two different cell lines, in the absence of cell cycle inhibitors (Haass et al., 2014; Spoerri et al., 2020). We then explore how we can model *intervention* experiments by using the mathematical model to replicate spheroid growth in the presence of a cell cycle inhibiting drug (Beaumont et al., 2016; Haass et al., 2014; Haass and Gabrielli, 2017; Vittadello et al., 2020).

### 4.1 Capturing intratumoral heterogeneity in a single cell line

Experimental images in Figures 2(a)-(c) illustrate the spheroid radius increases by a factor of approximately 1.3 over a period of 24 h. Over this timescale the intratumoural structure evolves from a relatively homogeneous mixture of G1 (red) and S/G2/M (green) cells at the beginning of the experiment to eventually form a complex structure comprising: (i) a necrotic core at the centre of the spheroid; (ii) an intermediate spherical shell containing live quiescent cells arrested in the G1 (red) phase; and (iii) an outer-most region composed of a mixture of freely proliferating G1 (red) and S/G2/M (green) cells. In our numerical explorations we seek to replicate these spatial and temporal patterns. For simplicity we always set *m*_1_ = *m*_2_ = *m*_3_ and *γ*(*c*) = 0, and assume that the initial spheroid is composed of an equal proportion of G2 (red) and S/G2/M (green) cells by setting *r*(*x,* 0) = *g*(*x,* 0) = 0.5. A MATLAB implementation of the numerical code is available on [GitHub](https://github.com/wang-jin-mathbio/jin2021) so that readers can explore the impact of choosing different parameter values.

First, we explore solutions of Equations (15)–(28) with the following carefully chosen parameter values: *C* = 1.8, *σ* = 1, *δ* = 0.9, *β* = 15, *c*_*r*_ = *c*_*g*_ = 0.1, *c*_*d*_ = 0.5, and *m*_1_ = *m*_2_ = *m*_3_ = 10. These choices give rise to the three kinetic rate functions shown in Figure 5(a). In this case *K*_*g*_(*c*) and *K*_*r*_(*c*) rapidly increase to their maximum rates, unity and *B* respectively, as *c* increases beyond the critical oxygen thresholds, *c*_*r*_ and *c*_*g*_, respectively. Similarly, *K*_*d*_(*c*) rapidly decreases to *C*(1 − *σ*) as *c* increases beyond the critical oxygen level *c*_*d*_. The visual depiction of the kinetic rate functions show that *K_r_*(*c*) > *K_g_*(*c*) for all *c*, so that the rate of G1 to S/G2/M transition is greater than the rate of S/G2/M to G1 transition. When oxygen is sufficiently reduced we have *K_d_*(*c*) > *K_r_*(*c*) > *K_g_*(*c*), which means that cell death is the dominant kinetic mechanism under these conditions. With these parameter choices we plot the evolution of *r*(*x, t*), *g*(*x, t*), *c*(*x, t*), *v*(*x, t*) and *l*(*t*) in Figure 6(a)-(c), where density and velocity profiles are plotted at *t* = 1.2, 1.8 and 2.4. These nondimensional times correspond to 12, 18, and 24 h, respectively, in the real experiment. Profiles in Figure 6(a) show that the spheroid grows with time and that both *g*(*x, t*) and *r*(*x, t*) are increasing functions of position, with the maximum volume fraction of living cells at the spheroid surface, *x* = *l*(*t*), and a minimum at the centre of the spheroid, *x* = 0. Profiles of *c*(*x, t*) also show that the oxygen concentration is an increasing function of *x*, which is consistent with oxygen diffusing into the spheroid from the free boundary at *x* = *l*(*t*), while simultaneously consumed by living cells. Comparing the profiles of *r*(*x, t*), *g*(*x, t*), and *c*(*x, t*), we see that by *t* = 2.4 we have almost a complete absence of living cells in the central region, indicating the eventual formation of a necrotic core, where *x* < 0.8.

**Figure 5:**
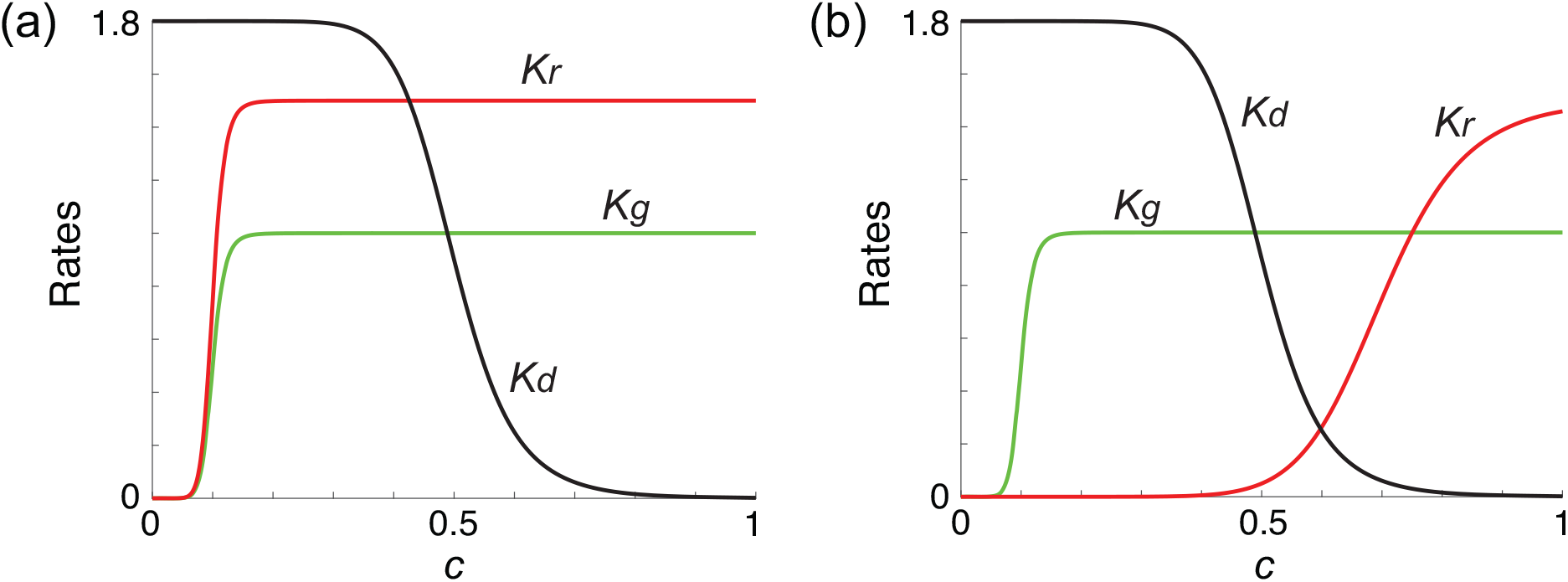
Kinetic rate functions *K_r_*(*c*), *K_g_*(*c*), and *K_d_*(*c*). (a) *c*_*r*_ = *c*_*g*_ = 0.1. (b) *c*_*r*_ = 0.7, *c*_*g*_ = 0.1, such that both *K_r_*(*c*) and *K_d_*(*c*) functions in (a) and (b) are identical. In both sets of profiles we have *B* = 1.5, *C* = 1.8, *c*_*d*_ = 0.5, *m*_1_ = *m*_2_ = *m*_3_ = 10.

**Figure 6:**
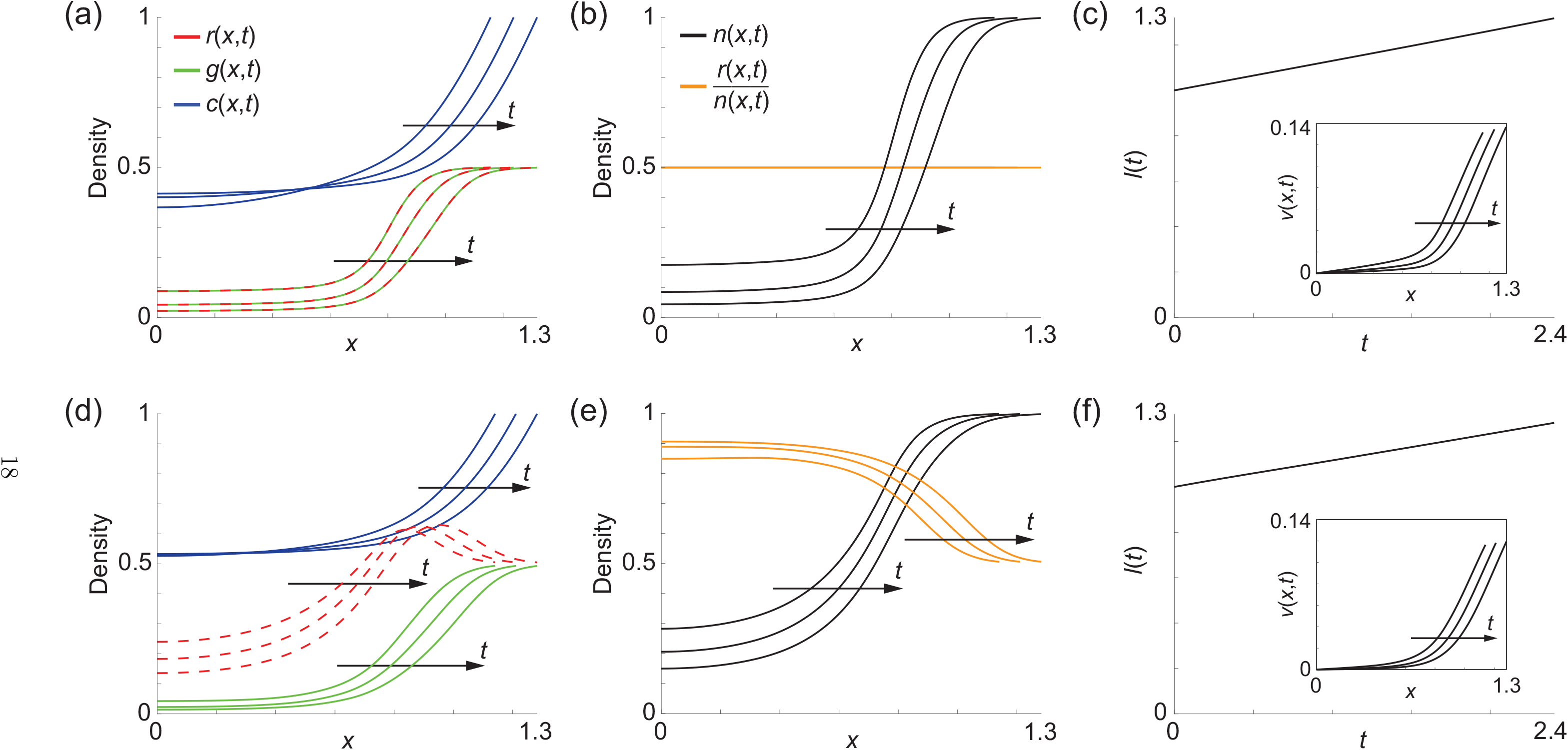
Numerical solutions of Equations (15)–(28) *t* = 1.2, 1.8, and 2.4. (a) and (d) Solutions *r*(*x, t*), *g*(*x, t*) and *c*(*x, t*), with the direction of increasing *t* indicated by the arrow. (b) and (e) Solutions *n*(*x, t*) = *r*(*x, t*) + *g*(*x, t*) and *r*(*x, t*)*/n*(*x, t*) as indicated. (c) and (f) Time evolution of spheroid radius, *l*(*t*), with the inset showing the velocity profile, *v*(*x, t*). Solutions in (a)-(d) correspond to the kinetic rate parameters in Figure 5(a), with *c*_*r*_ = *c*_*g*_ = 0.1. Solutions in (e)-(h) correspond to the kinetic rate parameters in Figure 5(b), with *c*_*r*_ = 0.7 and *c*_*g*_ = 0.1. In both simulations *B* = 1.5, *C* = 1.8, *σ* = 1, *δ* = 0.9, *β* = 15, *c*_*d*_ = 0.5, *m*_1_ = *m*_2_ = *m*_3_ = 10, *γ*(*c*) = 0.

An interesting feature of the solution in Figure 6(a) is that the spatial distribution of living cells involves two subpopulations where *r*(*x, t*) ≈ *g*(*x, t*) at all locations, at all times considered. Therefore, for this choice of parameters, while we do see the eventual formation of a necrotic core, we do not see the formation of an intermediate zone of G1-arrested cells like in Figure 2(c). This observation is clear when we plot the proportion of G1 cells, *r*(*x, t*)*/n*(*x, t*) in Figure 6(b), where we have *r*(*x, t*)*/n*(*x, t*) = 0.5 for all *x*, at all times considered. We visualise the cell velocity, *v*(*x, t*), in Figure 6(c) showing that we have *v*(*x, t*) > 0 for all *x* during the experiment, indicating that the total volume gain through proliferation exceeds the volume loss through cell death. This gain in volume leads to *l*(2.4) ≈ 1.3, which is consistent with the rate of expansion of the experimental spheroid in Figure 2(a)-(c) over the same time interval.

To emphasise the key features of this solution of Equations (15)–(28) we now summarise the solution from Figure 6(a) in a different format in Figure 7(a), by plotting *r*(*x, t*), *g*(*x, t*), *n*(*x, t*) and *c*(*x, t*) at the end of the experiment, *t* = 2.4, on *−l*(*t*) < *x* < *l*(*t*). Plotting the solution in this format helps to emphasise that the mathematical model can predict the formation of different intratumoral heterogeneity within the spheroid, as revealed by FUCCI labelling. We note that similar to Ward and King’s model, solutions of our model are continuous so that the internal boundaries between the three regions are not well-defined.

**Figure 7:**
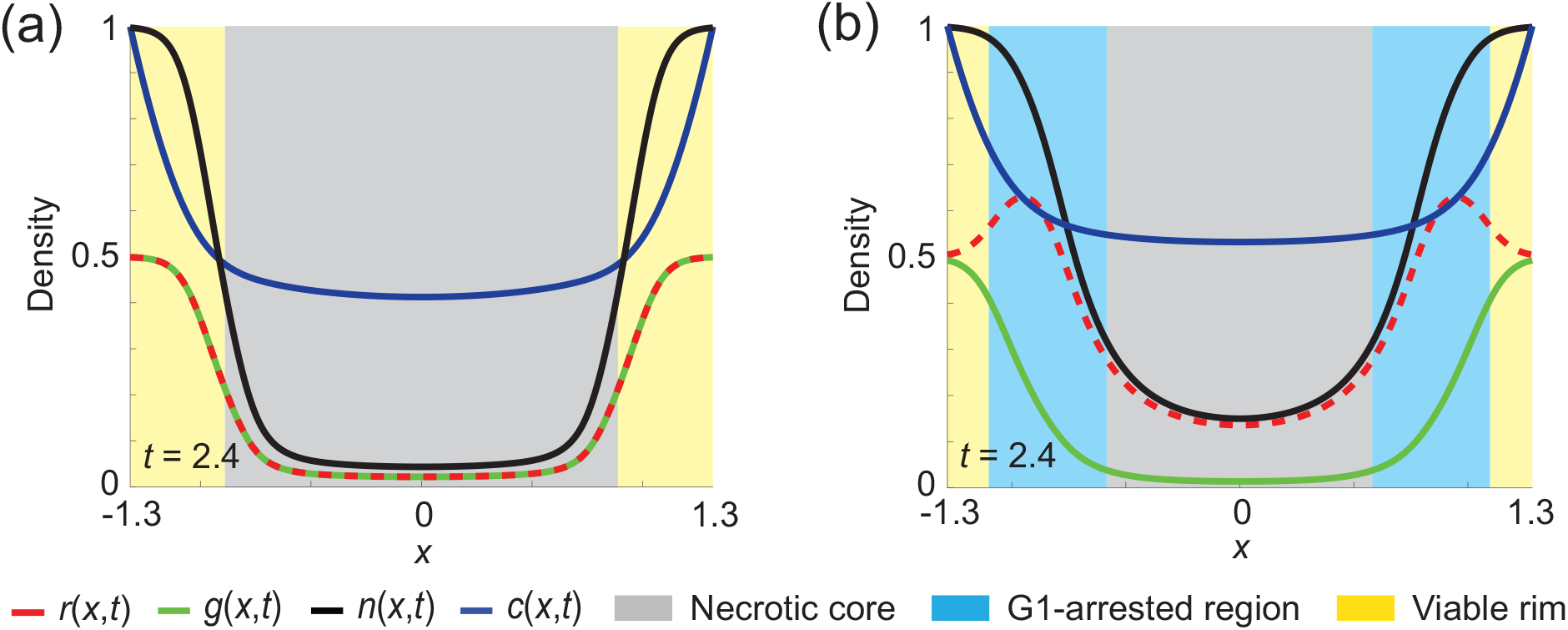
Visualisation of the predicted intratumoural structure. (a) and (b) show profiles of *r*(*x, t*), *g*(*x, t*), *n*(*x, t*), and *c*(*x, t*) at *t* = 2.4, on *l*(*t*) < *x* < *l*(*t*), with different intratumoural substructures highlighted by different colours. In particular we highlight the regions corresponding to the necrotic core, the intermediate G1-arrested quiescent zone and the freely-proliferating outer-rim. Results in (a) and (b) correspond to the parameter choice in Figure 6(a)-(c) and (d)-(f), respectively.

In Figure 7(a) we see that the distribution of *n*(*x, t*) and *g*(*x, t*) is such that an extensive central necrotic core develops, where |*x*| < 0.8, with very low density of living cells in the central region of the spheroid. In contrast, the outer region of the spheroid, where |*x*| > 0.8, is composed of freely proliferating cells with *g*(*x, t*) ≈ *r*(*x, t*). Within this outer-most regions there are no obvious spatial patterns of cell cycle status.

Our initial numerical results in Figure 6(a)-(c) are qualitatively consistent with the evolution of the spheroid size, *l*(*t*), measured in Figure 2. However, the intratumoural structure is very different since the numerical solution in Figure 6(a)-(c) does not involve any intermediate G1-arrested region as is clear in Figure 2(b)-(c). To incorporate this phenomenon, we resolve the model with *c*_*r*_ = 0.7 and *c*_*g*_ = 0.1, and all other parameters unchanged. The kinetic rate functions are shown in Figure 5(b) in which we see that: (i) *K*_*d*_(*c*) > *K*_*g*_(*c*) > *K*_*r*_(*c*) for sufficiently small oxygen concentrations; (ii) *K*_*g*_(*c*) > *K*_*d*_(*c*) > *K*_*r*_(*c*) at intermediate oxygen concentrations; and, (iii) *K*_*r*_(*c*) > *K*_*g*_(*c*) > *K*_*d*_(*c*) at sufficiently large oxygen concentrations. The associated numerical solutions of Equations (15)–(28) are shown in Figure 6(d)-(f). Comparing profiles in Figure 6(a) and Figure 6(d) indicates that we now have a very different intratumoural structure. Similar to before, we see that *g*(*x, t*) and *c*(*x, t*) are increasing functions of *x*, whereas *r*(*x, t*) is more complicated since there is a local maximum in *r*(*x, t*) within the spheroid. Interestingly, comparing the profiles of *n*(*x, t*) in Figure 6(b) and Figure 6(e) we see that the spatial distribution of the total live cell density is very similar for these two choices of parameters, yet the internal structure of the spheroids, as revealed by FUCCI-labelling in our theoretical model, is very different. The velocity profiles and the time evolution of the outer radius of the spheroid is shown in Figure 6(f), where again we see that the velocity field and the evolution of the outer spheroid radius is very similar for these two choices of model parameters.

The profile in Figure 7(b) highlights the intratumoural structure by plotting *r*(*x, t*), *g*(*x, t*), *n*(*x, t*) and *c*(*x, t*) at the end of the experiment, *t* = 2.4, on *−l*(*t*) < *x* < *l*(*t*). In this case we see that the internal structure defined by the distribution of various cell populations includes: (i) the necrotic core for |*x*| < 0.6; (ii) an intermediate region composed of G1-arrested (red) living cells for 0.6 < |*x*| < 1.1; and, (iii) an outer-most rim of freely proliferating cells for 1.1 < |*x*| < 1.3. This exercise of comparing results in Figure 6(a)–(c) and Figure 6(d)–(f) shows that the mathematical model we have developed can replicate key features of 3D tumour spheroid models with intratumoral heterogeneity revealed by FUCCI labelling. These differences are visually pronounced when we plot those results in Figure 7 where we deliberately highlight regions within the spheroid according to the proliferation status.

### 4.2 Capturing intratumoral heterogeneity in a multiple cell lines

Numerical results in Figure 6 and 7 show that our model is able to capture both the time evolution of the spheroid size, as well as the fundamental intratumoural structure regarding the status of the cell cycle at different positions within the spheroid for a single cell line, in this case the C8161 melanoma cell line. We now further explore the capacity of the model to represent different forms of intratumoural structure as revealed by FUCCI labelling in different cell lines. Very recent experimental data reported by Spoerri et al. (2020) describe a range of melanoma spheroids constructed using different cell lines, and characterise the proliferative heterogeneity calculating the percentage of G1-labelled (red) cells as a function of distance from spheroid’s surface. In our model we use the standard spherical coordinate system, 0 < *x* < *l*(*t*), so that *x* is the radial position measured relative to the centre of the spheroid. In Spoerri et al. (2020), we measure distance from the surface of the spheroid, *X* = *l*(*t*) − *x*, so that *X* = 0 is at the surface of the spheroid and *X* = *l*(*t*) is at the centre of the spheroid. Data reported by Spoerri et al. (2020) show the proportion of G1-labelled cells as a function of *X* gives rise to a range of different profiles that depend on the cell. Images in Figure 8(a)–(b) show melanoma spheroids with FUCCI labelling, using two different melanoma cell lines: WM164 cells in Figure 8(a) and C8161 cells in Figure 8(b). We show, schematically, the observed trends in terms of the proportion of G1-labelled cells as a function of *X* in Figure 8(c). For the WM164 cell line we see that the G1-labelled cells make up approximately 75 % of the total living population at the spheroid surface, and the proportion of G1-labelled cells increases gradually with *X*. In contrast, for the C8161 cell line, the proportion of G1-labelled cells is approximately 45 % of the total living population at the spheroid surface, and we see a far more rapid increase in the proportion of G1-labelled cells with *X*. These results are consistent with our visual interpretation of the spheroids in Figure 8(a)-(b) where we see that the WM164 spheroid does not contain a visually distinct necrotic core whereas the C8161 spheroid develops a very clear necrotic core. These differences are consistent with our previous measurements (Spoerri et al., 2020) showing that spatial differences in cell cycle status are more obvious in the C8161 spheroid relative to the WM164 spheroid.

**Figure 8:**
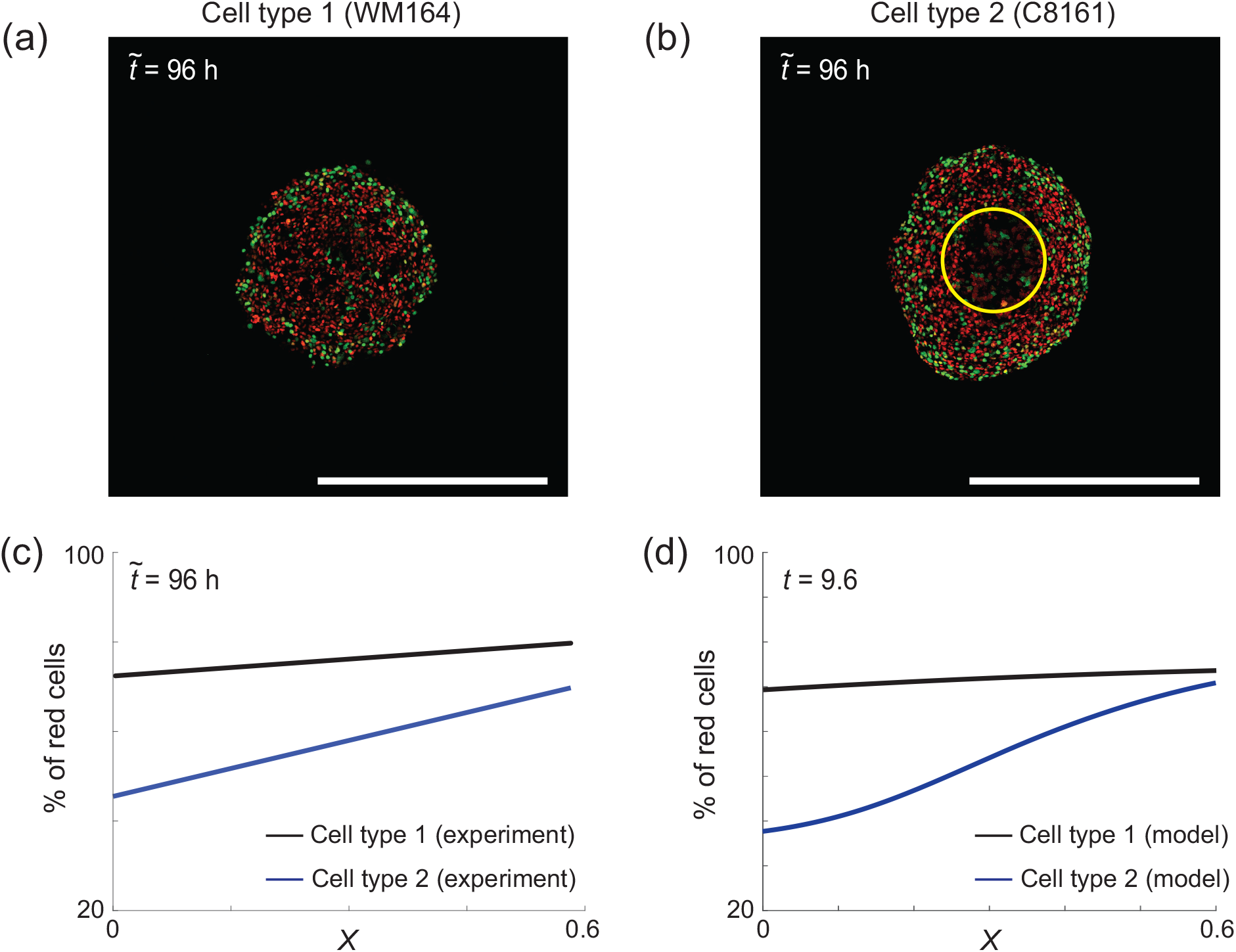
Comparison of experimental data and model predictions. (a) and (b) Confocal microscopy images of tumour spheroids after 96 h derived from two different cell types. The scale bar corresponds to 400 *μ*m. (c) Schematics showing different profiles of the percentage of red cells within the spheroids. (d) Numerical results qualitatively reproducing experimental data shown in (c). For the simulation of cell type 1 (WM164), *B* = 1, *c*_*r*_ = 0.9, *m*_1_ = *m*_2_ = *m*_3_ = 3. For the simulation of cell type 2 (C8161), *B* = 3, *c*_*r*_ = 0.7, *m*_1_ = *m*_2_ = *m*_3_ = 6. For both simulations, *C* = 1.8, *σ* = 1, *δ* = 0.9, *β* = 10, *c*_*g*_ = 0.1, *c*_*d*_ = 0.5, *γ*(*c*) = 0.

Results in 8(d) show numerical solutions of Equations (15)–(28) with carefully chosen parameter values. In particular we choose *B* = 1, *c*_*r*_ = 0.9, and *m*_1_ = *m*_2_ = *m*_3_ = 3 for the WM164 spheroid, and *B* = 3, *c*_*r*_ = 0.7, and *m*_1_ = *m*_2_ = *m*_3_ = 6 for the C8161 spheroid. With these parameter values we obtain the solution of Equations (15)–(28) at *t* = 9.6, and we plot 100 *× r*(*X, t*)*/n*(*X, t*) %, revealing that the mathematical model is sufficiently flexible to capture these kinds of intratumoral heterogeneity reported by Spoerri et al. (2020).

### 4.3 Simulating the action of anti-mitotic drugs

In Sections 4.1–4.2 we focus on using the mathematical model to replicate and visualise spatial patterns of cell cycle status within a growing spheroid under control conditions. One of the key reasons that 3D spheroid cultures are of high interest in the experimental cancer cell research community is the ability to test different potential anti-cancer drug treatments in a realistic 3D geometry (Friedrich et al., 2009; Loessner et al., 2013). Various anti-mitotic drugs have been developed and applied to 3D spheroid cultures (e.g. Crivelli et al., 2012; Loessner et al., 2013; Smalley et al. 2006), and with the development of FUCCI-labelling we have vastly improved opportunities to visualise how such drug treatments impact the spatial distribution of cell cycle status within the treated spheroid (Haass et al., 2014; Beaumont et al., 2016; Kienzle et al., 2017). The experimental images in Figure 9(a) and Figure 9(d) show a control C8161 melanoma spheroid, and a U0126-treated C8161 melanoma spheroid, after six days, respectively. In this case, both the control and treatment spheroids are grown for three days initially, and U0126 is applied at day three to the treatment group. U0126 is a MEK-inhibitor that leads to G1 arrest (Haass et al., 2014; Smalley et al. 2006). Comparing the images in Figure 9(a) and Figure 9(d) leads to two obvious conclusions: (i) the treated spheroid is almost completely composed of G1-arrested (red) cells; and, (ii) the size of the treated spheroid is smaller than the untreated spheroid.

**Figure 9:**
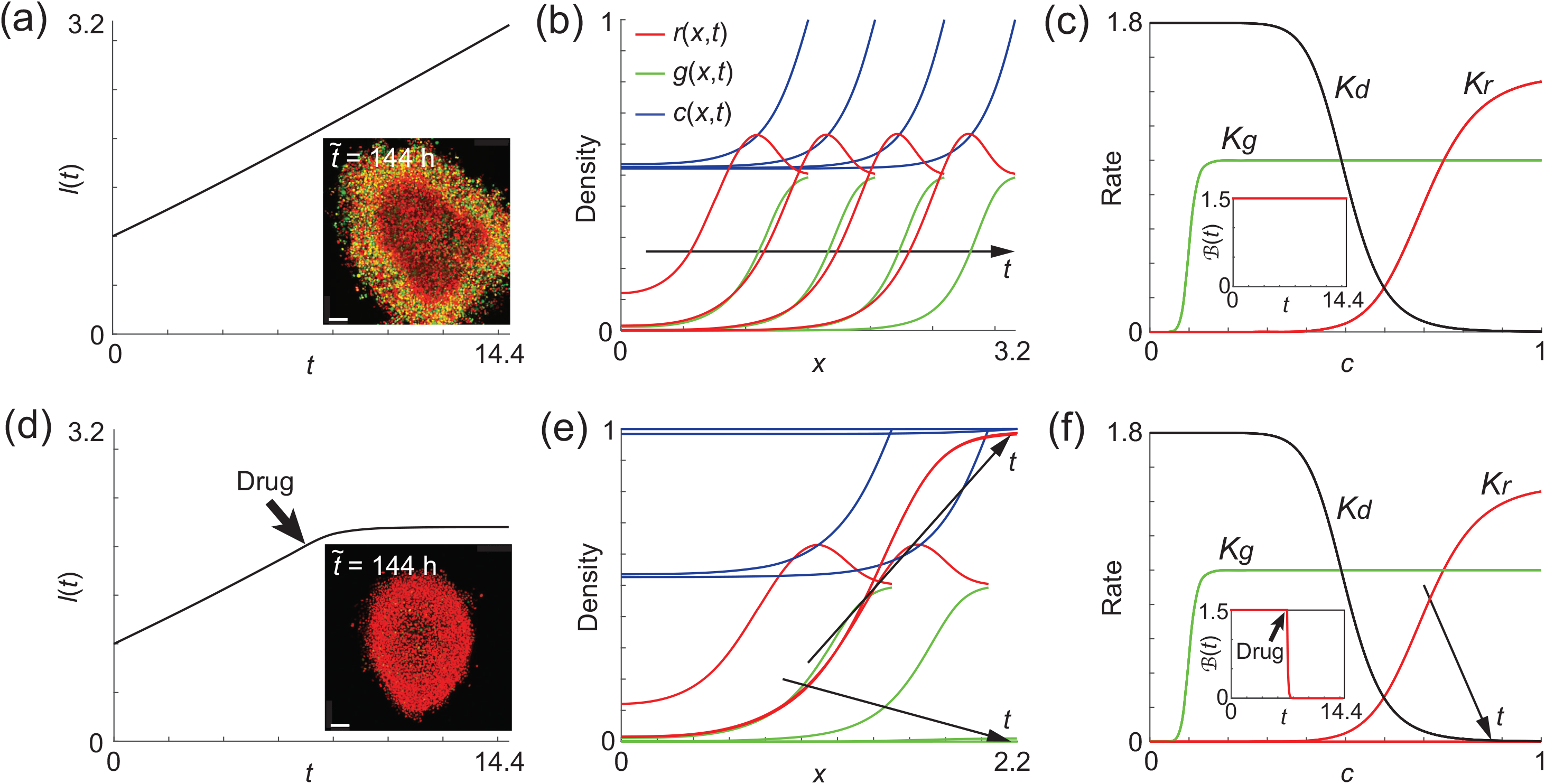
Numerical results showing the effects of drug on the heterogeneity of cell phases within a tumour spheroid. (a)-(c) Profiles of tumour spheroid growth without treatment (control). The scale bar corresponds to 110 *μ*m. (d)-(f) Profiles of tumour spheroid growth with the drug applied at *t* = 7.2. (a) and (d) Time evolution of the radius with and without treatment, respectively. The insets show the related experimental images of melanoma spheroids at day six, with the scale car corresponding to 110 *μ*m. (b) and (e) Density profiles of *r*, *g*, and *c* at *t* = 3.6, 7.2, 10.8 and 14.4. (c) and (f) Profiles of cell kinetics. In both simulations, *C* = 1.8, *σ* = 1, *δ* = 0.9, *β* = 10, *c*_*r*_ = 0.7, *c*_*g*_ = 0.1, *c*_*d*_ = 0.5, *m*_1_ = *m*_2_ = *m*_3_ = 10, *γ*(*c*) = 0.

To incorporate the effect U0126 in our model we make the straightforward assumption that the drug affects the maximum rate at which cells transition from G1 phase (red) to S/G2/M phase (green) by modifying the appropriate kinetic function,

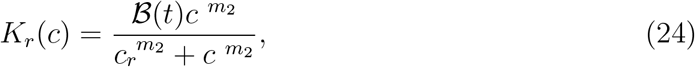

with

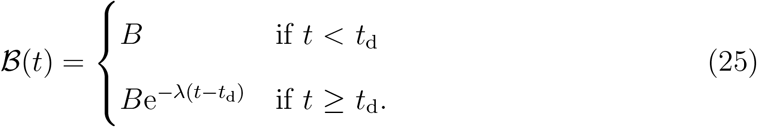

Here, *t*_d_ is the time that the drug is applied, *λ* > 0 is an intrinsic drug-sensitivity parameter (Enderling and Chaplain, 2014; Lewin et al., 2018). We note that the choice of the form of drugs is not unique, and can vary depending on the type of cell and treatment (Collis et al., 2017; Crivelli et al., 2012; de Pillis et al., 2006).

We simulate these experiments by setting *t*_d_ = 7.2 and *λ* = 10, and summarise our numerical results in Figure 9. Comparing profiles in Figure 9(a) and Figure 9(d) we see that the control and treatment experiments are indistinguishable for *t < t*_d_, as expected. For *t > t*_d_ we see a clear reduction in the time rate of change of radius, *l*(*t*), in the treated spheroid. The treated tumour has almost stopped growing at the end of the simulation whereas the untreated spheroid is continues to grow at an almost constant rate of radial expansion. The density profiles in Figure 9(b) show that the control spheroid rapidly develops a necrotic core, and that the populations of live cells continues to expand over time. In comparison, the density profiles in Figure 9(e) show that after the application of the drug we observe a rapid decline in the *g*(*x, t*) subpopulation and the total population becomes almost entirely composed of the *r*(*x, t*) subpopulation. Note that the density profiles in Figure 9(b) and Figure 9(e) are plotted on different spatial scales so that the intraturmoral structure is visually clear. The associated kinetic rate functions for these simulations are summarised in Figure 9(c) and Figure 9(f), with the drug treatment summarised by the function 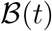, shown as an inset in both subfigures.

## 5 Conclusion and Outlook

In this work we develop a new mathematical model of 3D tumour spheroid growth that is compatible with FUCCI-labelling, which highlights the spatial location of cells in terms of their cell cycle status. In these experiments cells in G1 phase fluoresce red, while cells in S/G2/M phase fluoresce green. Traditional tumour spheroid experiments do not provide spatio-temporal information on cell cycle status, and it is not obvious whether all living cells are freely cycling or not. Our experiments show that FUCCI provides detailed insight into the development of intratumoral heterogeneity as the spheroid grows. In particular, at late time points we see: (i) a necrotic region in the centre of the spheroid that does not contain living fluorescent cells; (ii) an intermediate spherical shell composed of living G1-arrested (red) cells; and, (iii) an outer spherical shell composed of a mixture of freely cycling G1 (red) and S/G2/M (green) cells. We demonstrate how to incorporate this information into a model of 3D tumour spheroids by adapting the well–known model of Ward and King (1997) to include three subpopulations of cells: (i) cells in G1 phase, *r*(*x, t*); (ii) cells in G2/S/M phase, *g*(*x, t*), and (iii), dead cells *m*(*x, t*). In this framework we assume cells are transported by the local velocity field, and the transitions between the G1 and G2/S/M phases are consistent with FUCCI-labelling. Rates of cell death and time points of transition through the cell cycle are taken as functions of the local oxygen concentration. Further, we consider oxygen to diffuse into the spheroid from the free surface, while being simultaneously consumed by the living cells. We show how to partially parameterise the mathematical model using experimental observations, and we explore numerical solutions of the nonlinear moving boundary problem and show that this model can capture key features of intratumoral heterogeneity in a single cell line, differences in intratumoral heterogeneity across multiple cell lines, as well as being able to mimic spheroid growth in the presence of anti-mitotic drugs.

There are many opportunities for further generalisation of this work, both experimentally and theoretically. From a modelling perspective, one of the key assumptions we make in modelling FUCCI is that we consider two live subpopulations of cells: G1 phase (red), and S/G2/M phase (green). In reality, it is also possible to identify an additional subpopulation where both the red and green fluorescence are active, giving rise to a yellow subpopulation that is often considered to be in early S phase (Haass et al., 2014). Under these conditions we would consider the living population to be composed of three subpopulations: G1 phase (red), eS phase (yellow) and S/G2/M phase (green). Extending our model to deal with this additional phase would involve reformulating the mathematical model to describe three subpopulations of living cells and one subpopulation of dead cells. This extended model would be able to capture additional biological features, but this benefit comes with the cost of incorporating a greater number of model parameters need to be estimated. While the mathematical extension to deal with eS (yellow) is relatively straightforward, the parameterisation and interpretation of this extended model is less obvious, and so we leave this extension for future consideration. Another opportunity for further investigation is the analysis of travelling wave solutions of Equations (15)–(28). In their original work, Ward and King (1997) were able to make progress in interpreting their model in terms of travelling wave analysis in the limit that the exponents in the Hill function become infinitely large and the kinetic rate functions simplify to Heaviside functions. We anticipate that a similar analysis could be considered for our more complicated model, and we leave this analysis for future consideration. From an experimental perspective we note that all experimental images and data discussed in this work relate to tumour spheroids composed of melanoma cells. However, we anticipate that our model and modelling framework will also be relevant to spheroids made from other types of cancer cell lines transduced with FUCCI system. As already noted, our experimental motivation involves the traditional FUCCI system with two fluorescent colours, which motivates us to consider the living cells as being composed of two subpopulations. More recent experimental technology, FUCCI4 (Bajar et al., 2016) uses four florescent colours to label the cell cycle. We expect that our approach here could be developed further by considering the live cell population to be composed of four subpopulations to model tumour spheroids with FUCCI4 labelling.

## APPENDIX A Ward and King (1997)

Since the new mathematical model developed in this work is an extension of Ward and King’s model (1997), we begin by presenting a robust numerical method to solve the original partial differential equation (PDE) model. In this preliminary Section we consider the tumour spheroid to be composed of two subpopulations: (i) living cells with volume fraction *n*(*x, t*), and (ii) dead cells with volume fraction *m*(*x, t*). The nondimensional PDE system is given by

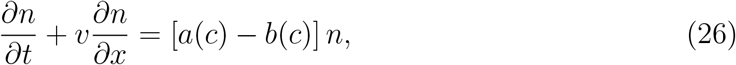

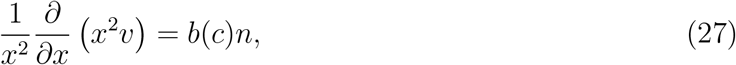

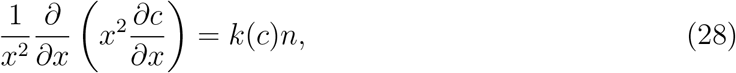

on 0 < *x* < *l*(*t*) and *t* > 0. Here, *v*(*x, t*) is the velocity field, *c*(*x, t*) is the oxygen concentration, *a*(*c*) = *k_m_*(*c*) *−k_d_*(*c*), *b*(*c*) = *k_m_*(*c*) *−*(1 *−δ*)*k*_*d*_(*c*), and *k*(*c*) = *βk_m_*(*c*)+*γ*(*c*). The original dimensional model, together with the rescaling arguments are outlined in Ward and King (1997).

The nondimensional kinetic rate functions, *k*_*m*_(*c*) and *k*_*d*_(*c*), are given by

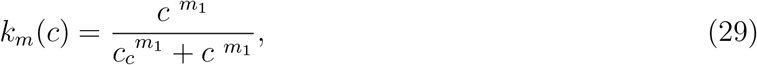

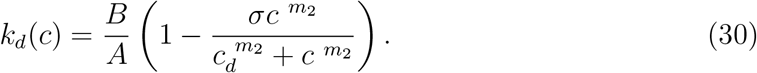

The initial and boundary conditions are

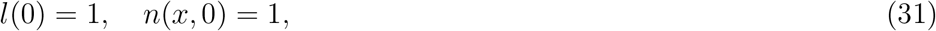

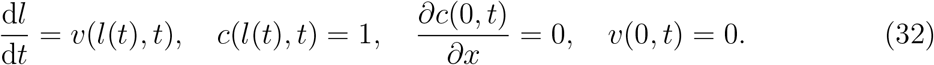

## APPENDIX B Numerical method for Ward and King (1997)

We solve Equations (26)–(28) by introducing a boundary fixing transformation *x* = *l*(*t*)*ξ* (Fadai and Simpson, 2020), so that the model becomes

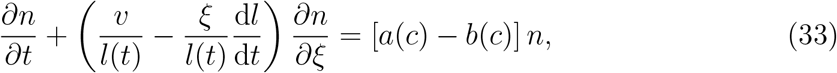

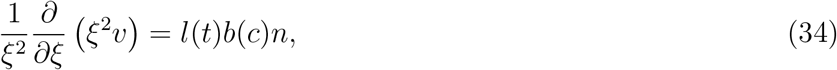

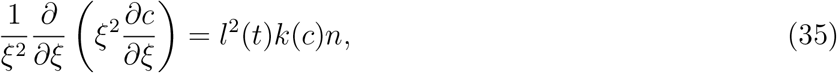

on 0 < *ξ* < 1. The transformed boundary conditions are

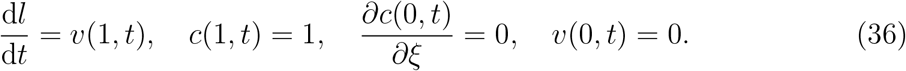

We discretise Equations (33)–(35) on a uniform mesh with spacing *δξ*. Central differences are used to approximate the terms associated with diffusive transport (Simpson et al., 2005), and terms associated with advection are approximated using upwinding,

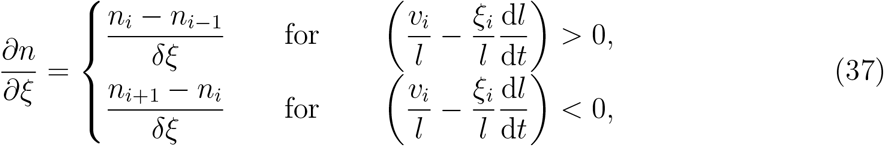

where 1 ≤ *i* ≤ *N* is the index for spatial nodes. Note that for the moving boundary we have d*l/*d*t* = *v*(*l*(*t*), *t*). Temporal derivatives in Equations (33)–(35) are approximated using a backward Euler approximation with time step *δt* (Morton and Mayers, 2005). The resulting system of nonlinear algebraic equations are solved using the Newton-Raphson method with convergence tolerance ∊. After the Newton-Raphson iterations converge, we update *l*(*t*). Software written in MATLAB are available on [GitHub](https://github.com/wang-jin-mathbio/jin2021). Results in Figure 10 show profiles that reproduce results from Figures 1–4 in Ward and King (1997).

**Figure 10:**
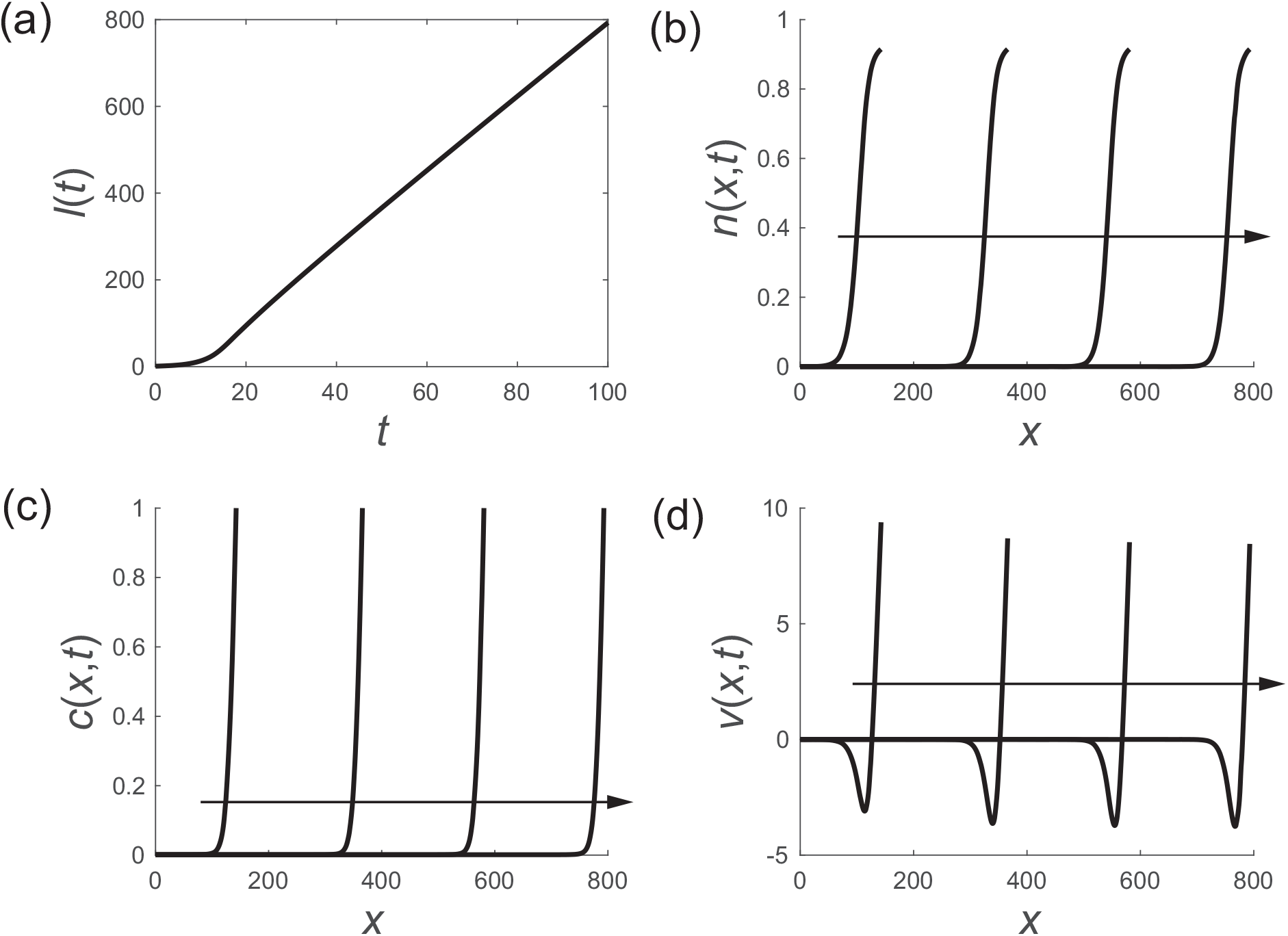
Numerical results for Ward and King’s model at *t* = 25, 50, 75, 100. (a) Time evolution of spheroid radius. (b) Profiles of living cell density. (c) Profiles of oxygen concentration. (d) Profiles of cell velocity. The arrow indicates increasing time. *B/A* = 1, *σ* = 0.9, *δ* = 0.5, *β* = 0.005, *c_c_* = 0.1, *c*_*d*_ = 0.05, *m*_1_ = *m*_2_ = 1.

## APPENDIX C Numerical method for model describing tumour spheroids with FUCCI labelling

To solve Equations (15)–(18) in the main document we introduce a boundary fixing transformation *x* = *l*(*t*)*ξ* to give

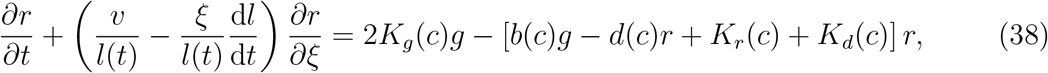

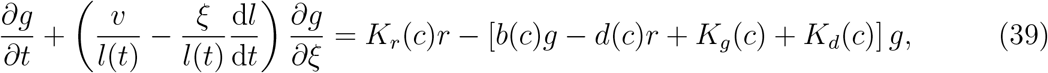

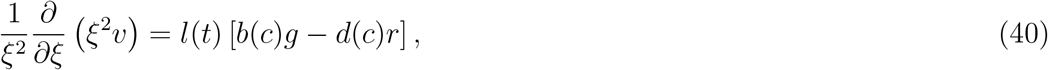

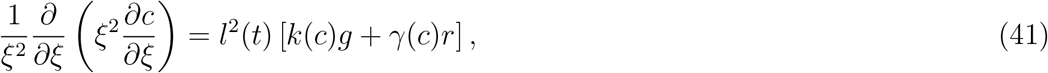

on 0 < *ξ* < 1. The transformed boundary conditions are

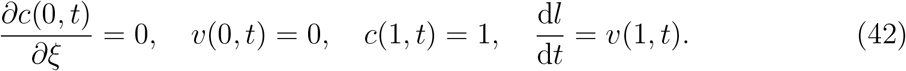

Similar to the numerical method outlined in Section 5, we discretize Equations (38)–(40) on a uniform mesh with spacing *δξ*. Central differences are used to approximate the terms arising from the diffusive transport terms, and upwinding is used to approximate the terms associated with advective transport. Temporal derivatives are approximated using a backward Euler approximation with a uniform time step *δt*. Newton-Raphson iterations are used to solve the resulting system of nonlinear algebraic equations with convergence tolerance *E*. After the Newton-Raphson iterates converge, we update *l*(*t*) using the boundary condition d*l/*d*t* = *v*(1, *t*). For all numerical simulations in this work we choose *δt* = *δξ* = 1 *×* 10^−3^ and *E* = 1 *×* 10^−6^, which leads to grid-independent results for the parameter values that we consider. Software written in MATLAB are available on [GitHub](https://github.com/wang-jin-mathbio/jin2021).

## APPENDIX D Sensitivity analysis

Here we study the sensitivity of the spheroid size, *l*(*t*), in terms of the model parameters. In the nondimensional system we have 11 free parameters, and were we focus on the parameter values associated with the results for cell type 2 in Figure 8(b) and Figure 8(d). In this case we have (*δ, β, σ, B, C, c_g_, c_r_, c_d_, m*_1_, *m*_2_, *m*_3_) = (0.9, 10, 1, 3, 1.8, 0.1, 0.7, 0.5, 6, 6, 6). To examine the sensitivity of the model solution to these choices we compute *l*(2.4) for the base set of parameters, and then systematically re–compute *l*(2.4) by varying each parameter, one at a time, by ±10%, keeping the other parameters constant.

Results in Figure 11 show *l*(2.4) = 1.365 as a horizontal dashed line. Predictions of *l*(2.4) when each parameter is varied in each column of Figure 11. For example, in the first column we examine the sensitivity of *l*(2.4) to the choice of *δ*, with the red and blue crosses indicating *l*(2.4) when *δ* is increased and decreased by 10%, respectively. Note that this combination of parameters has *σ* = 1, which is the maximum value possible. Therefore, for the sensitivity of *σ*, we only show *l*(2.4) when the parameter is reduced by 10%. In summary, the sensitivity results in Figure 11 indicate that certain parameters, such as *c*_*r*_, and *c*_*d*_, play a key role since *l*(2.4) is most sensitive to these parameters. Note that this simple parameter sensitivity analysis provides important initial insight, however we leave a more thorough sensitivity analysis for future consideration.

**Figure 11:**
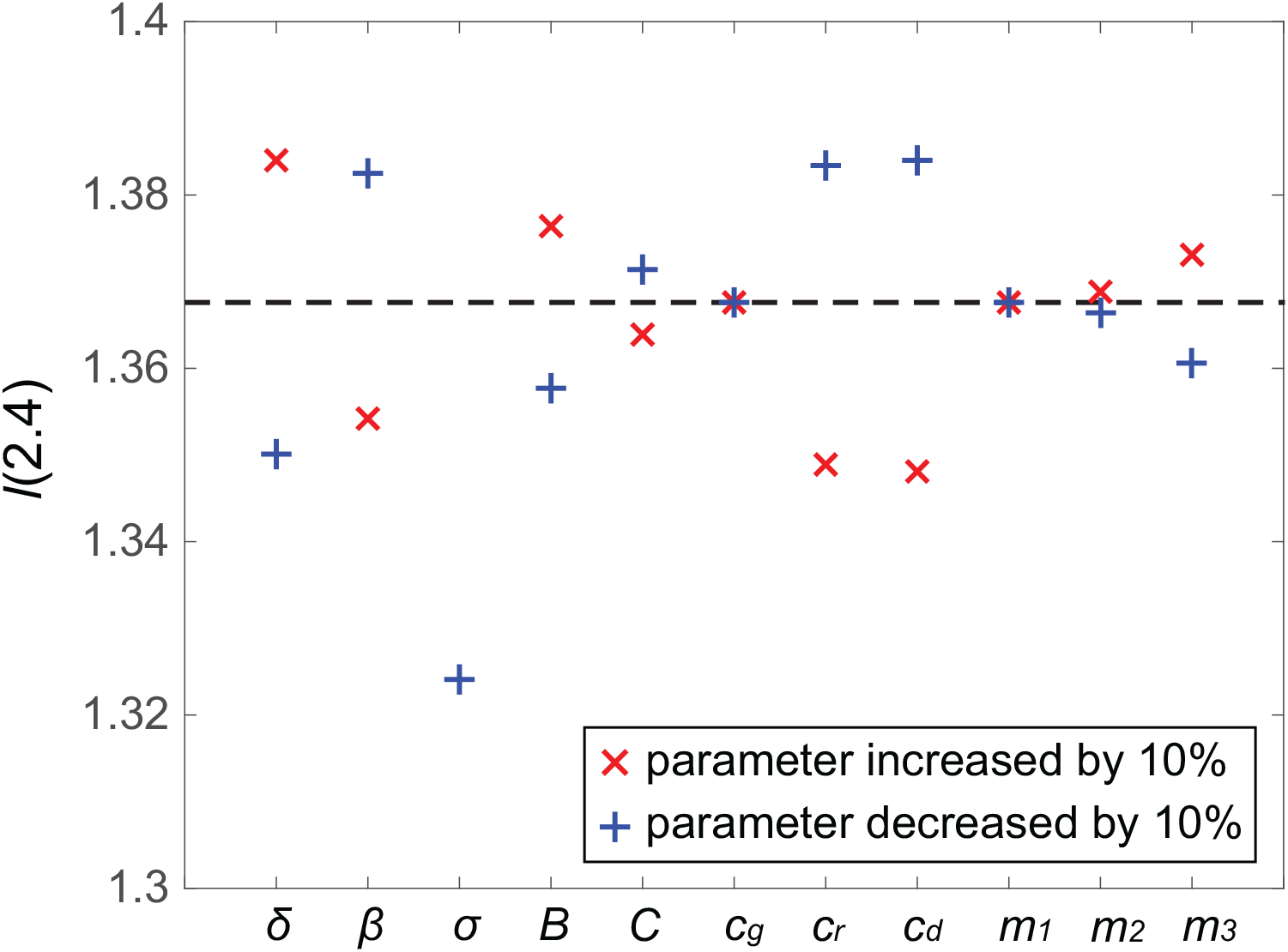
Numerical results of sensitivity analysis. The dashed line indicates the benchmark value *l*(2.4) = 1.365. The red and blue markers vertically aligned with each parameter indicate the value of *l*(2.4) when each parameter is varied by ±10%, respectively. Note that there is no red marker for *σ* due to that *σ* = 1 is the largest value this parameter can take.

## Acknowledgements

MJS and NKH are supported by the Australian Research Council (DP200100177). NKH is supported by the National Health and Medical Research Council (APP1084893). WJ is supported by a QUT Vice-Chancellor’s Research Fellow-ship.

